# Leaf Variegation and Impaired Chloroplast Development Caused by a Truncated CCT Domain gene in *albostrians* Barley

**DOI:** 10.1101/560797

**Authors:** Mingjiu Li, Goetz Hensel, Martin Mascher, Michael Melzer, Nagaveni Budhagatapalli, Twan Rutten, Axel Himmelbach, Sebastian Beier, Viktor Korzun, Jochen Kumlehn, Thomas Börner, Nils Stein

## Abstract

Chloroplasts fuel plant development and growth by converting solar into chemical energy. They mature from proplastids through the concerted action of genes in both the organellar and the nuclear genome. Defects in such genes impair chloroplast development and may lead to pigment-deficient seedlings or seedlings with variegated leaves. Such mutants are instrumental as tools for dissecting genetic factors underlying the mechanisms involved in chloroplast biogenesis. Characterization of the green-white variegated *albostrians* mutant of barley has greatly broadened the field of chloroplast biology including the discovery of retrograde signaling. Here, we report the identification of the *ALBOSTRIANS* gene *HvAST* by positional cloning as well as its functional validation based on independently induced mutants by TILLING and RNA-guided Cas9 endonuclease mediated gene editing. The phenotypes of the independent *HvAST* mutants imply residual activity of HvAST in the original *albostrians* allele conferring an imperfect penetrance of the variegated phenotype even at homozygous state of the mutation. *HvAST* is a homolog of the *Arabidopsis thaliana CCT Motif* transcription factor gene *AtCIA2*, which was reported to be involved in the expression of nuclear genes essential for chloroplast biogenesis. Interestingly, in barley we localized HvAST to the chloroplast indicating novel without any clear evidence of nuclear localization.

**One-sentence summary:** Leaf variegation in the barley mutant *albostrians* is caused by mutation of a single CCT-domain containing gene with residual activity, which is directed to the chloroplast.

## INTRODUCTION

Chloroplasts are the site of photosynthesis. In higher plants, functional chloroplasts differentiate from their progenitor organelles called proplastids. This process, chloroplast biogenesis, depends on a network of environmental, temporal and cellular factors (Pogson and Albrecht, 2011), with the latter including the expression of both plastid and nuclear genes. The nuclear genome codes for the vast majority of proteins required for chloroplast biogenesis. In barley, for instance, only 78 out of about 3000 chloroplast proteins are encoded in the plastid genome (plastome) (Saski et al., 2007; Petersen et al., 2013). Mutations in these nuclear genes may result in defective plastids as reflected by frequently occurring leaf coloration aberrations in different plant species. Chlorophyll-deficient mutants provide valuable genetic tools for the identification of nuclear genes involved in chloroplast biogenesis and regulation (Taylor et al., 1987; Barkan, 1998; Leon et al., 1998).

Specifically, mutations leading to variegation, due to their unique feature of exhibiting both normal and defective plastids in different sectors of the same tissue, are of great interest to research towards understanding (i) chloroplast biogenesis, (ii) cross-communication between the nucleus and the other DNA-containing compartments like plastids and mitochondria, (iii) and the molecular mechanism of variegation. Genes involved in the variegation pattern formation have been cloned in monocot and dicot species (e.g. Wu et al., 1999; Chen et al., 2000; Takechi et al., 2000; Prikryl et al., 2008; Hayashi-Tsugane et al., 2014; Wang et al., 2016; Zheng et al., 2016; Guan et al., 2017; Zagari et al., 2017). Despite the advances of identifying the underlying genes, insights to the molecular mechanisms controlling the phenomenon are rare (Sakamoto, 2003; Yu et al., 2007). Leaf variegation can occur when cells of green and pigment-deficient sectors have distinct genotypes. Cells in green sectors have a wild-type genotype with functional chloroplasts, while pigment-deficient sectors consist of mutant cells with abnormal plastids. Multiple mechanisms, such as somatic chimerism, transposable elements activity, and organellar genome mutations, are involved in generating cells of a single plant with distinct genotypes (Yu et al., 2007). In other cases, variegation occurs also in mutants with identical genotype in the green and the chlorotic tissue sectors. Especially the latter cases provide interesting genetic systems for investigations into the poorly understood pathways of chloroplast biogenesis (Sakamoto, 2003; Putarjunan et al., 2013).

Due to the differences in leaf organization between monocots and dicots, ‘variegation’ in monocots is also referred to as ‘striping’. The *iojap* and *albostrians* mutants of maize and barley, respectively, are two well-studied examples of mutations causing a striping phenotype. They belong to the group of mutants with identical genotype in green and pigment-deficient tissues. The *iojap* gene codes for a component of the 50S subunit of the plastid ribosome. Its role in ribosome assembly or function is yet unknown (Han et al., 1992; Wanschers et al., 2012). The *albostrians* mutant served as a model for studies on the transcriptional machinery of the chloroplast genome (Hess et al., 1993; Zhelyazkova et al., 2012) and on regulatory interactions between plastids, mitochondria and the nucleus (Bradbeer et al., 1979; Hess et al., 1994; Hedtke et al., 1999; Nott et al., 2006). However, elucidation of the mechanism leading to the *albostrians*-specific phenotype of variegation was impeded by the fact that the causal underlying gene was unknown. The *albostrians* mutant was selected in the 1950s after X-ray irradiation of the two-rowed spring barley variety ‘Haisa’. The phenotype is controlled by a single recessive nuclear gene, which, however, has no complete penetrance; the progeny of the fully green mutant segregates into green, variegated and albino seedlings in a ratio of around 1:8:1 (Hagemann and Scholz, 1962). Green and variegated seedlings with sufficient photosynthetically active green tissue will grow to maturity, remain green and/or striped, respectively, and produce fertile flowers. The progeny again will consist of green, white and variegated plants (Hagemann and Scholz, 1962). The phenotype is very stable since changes of light conditions or temperature have remained without effect (Hagemann and Scholz, 1962; Börner et al., 1976; Hess et al., 1993). Similar to *iojap* (Walbot and Coe, 1979), the barley *HvAST* allele [termed *Hvas* in (Hagemann and Scholz, 1962)] was originally described as a ‘plastome mutator’, *i.e.* as a nuclear gene that induces plastome mutations because the nuclear-gene induced albino phenotype showed a stable maternal inheritance (Hagemann and Scholz, 1962). This hypothesis was underpinned originally by the finding of so-called ‘mixed cells’ containing undifferentiated plastids and green chloroplasts (Knoth and Hagemann, 1977). Later, it has been proposed, however, that neither *iojap* nor *HvAST* are mutator genes since in either mutant the albinotic leaf tissues were characterized by ribosome-deficient plastids (Börner et al., 1976; Walbot and Coe, 1979; Börner and Hess, 1993) and the lack of plastid ribosomes is stably inherited like a genuine plastome mutation (Zubko and Day, 1998). Indeed, restriction patterns and sequence of the DNA in undifferentiated, ribosome-free plastids of *albostrians* were identical with those of the chloroplast DNA in green tissues of *albostrians* and in the barley wild-type varieties ‘Haisa’ and ‘Morex’ (Hess et al., 1993; Zhelyazkova et al., 2012). Thus, the barley *HvAST* gene product is not expected to act on chloroplast DNA but to be required for plastid ribosome assembly, maintenance and/or function.

Recently, ample genomic resources have been developed in barley (Schulte et al., 2009; Mayer et al., 2011; International Barley Genome Sequencing Consortium, 2012; Ariyadasa et al., 2014; Mascher et al., 2017) which are greatly facilitating gene cloning (Mascher et al., 2014). Using these resources, we identified a candidate gene by positional cloning and confirmed its identity with *ALBOSTRIANS* by screening of a barley TILLING population (Gottwald et al., 2009) and by site-directed mutagenesis using RNA-guided Cas9 endonuclease (Jinek et al., 2012). The gene *HvAST* underlying the *albostrians* phenotype of variegation is identical with *HvCMF7*, a member of the *CCT Motif* gene *Family* (*CMF*) of putative transcription factors (Cockram et al., 2012). The transient analysis of an HvAST:GFP fusion in conjunction with organelle markers revealed a strong subcellular localization of ALBOSTRIANS to the chloroplast.

## RESULTS

### Map-based Cloning of the *HvAST* Gene

The mutant *Hvast* allele of barley causes variegation, characterized by green-white-striped leaves (Figure 1A). The white leaf sectors contain undifferentiated plastids virtually free of 70S ribosomes (Hess et al., 1993) while green parts harbor normal, photosynthetically active chloroplasts. We allocated the gene *HvAST* to the long arm of chromosome 7H by using two F_2_ mapping populations [Morex x M4205 (MM4205) and Barke x M4205 (BM4205)] each representing 182 gametes. Further fine mapping of the gene in 2,688 gametes of population MM4205 delimited the *HvAST* target region to a 0.06 cM interval (Figure 1B). The closest *HvAST* flanking markers were anchored to the physical map of barley (International Barley Genome Sequencing Consortium, 2012; Mascher et al., 2013) and by sequence comparison and BAC library screening, five overlapping physical map contigs (Finger Printed Contigs, FPcontigs) spanning a distance of around 0.40 Mbp (between flanking markers Zip_2661 and 3_0168) were identified. Then, sixty BAC clones representing the physical target interval were sequenced providing novel information for developing and mapping of new BAC contig derived markers; delimiting the *HvAST* locus to two overlapping BAC clones HVVMRXALLrA0395M21 and HVVMRXALLmA0230A06 (Figure 1C). Eventually, a 46 Kbp region was found to be the smallest genetic and physical interval comprising the *HvAST* locus (Figure 1D). Sequence comparison to annotated genes defined on the Morex draft WGS assembly (International Barley Genome Sequencing Consortium, 2012) revealed the presence of a single gene locus, *MLOC_670* (GenBank accession ID AK366098). The *HvAST* gene structure, supported by cDNA analysis, consists of three instead of two exons as *in silico* predicted for *MLOC_670*. We discovered a 4 bp deletion by re-sequencing of the *HvAST* gene in the *albostrians* mutant (line M4205) if compared to the wild-type genotype (Figure 1E). This mutation is predicted to induce a shift of reading frame and, as a consequence, a premature stop codon in the second exon of the gene. This lesion is expected to result in (partial) loss of the gene function in the mutant and makes the *MLOC_670* gene locus a very strong candidate for representing the *HvAST* gene. The mutant allele in the original *albostrians* mutant is designated as *Hvast1*.

**Figure 1.**
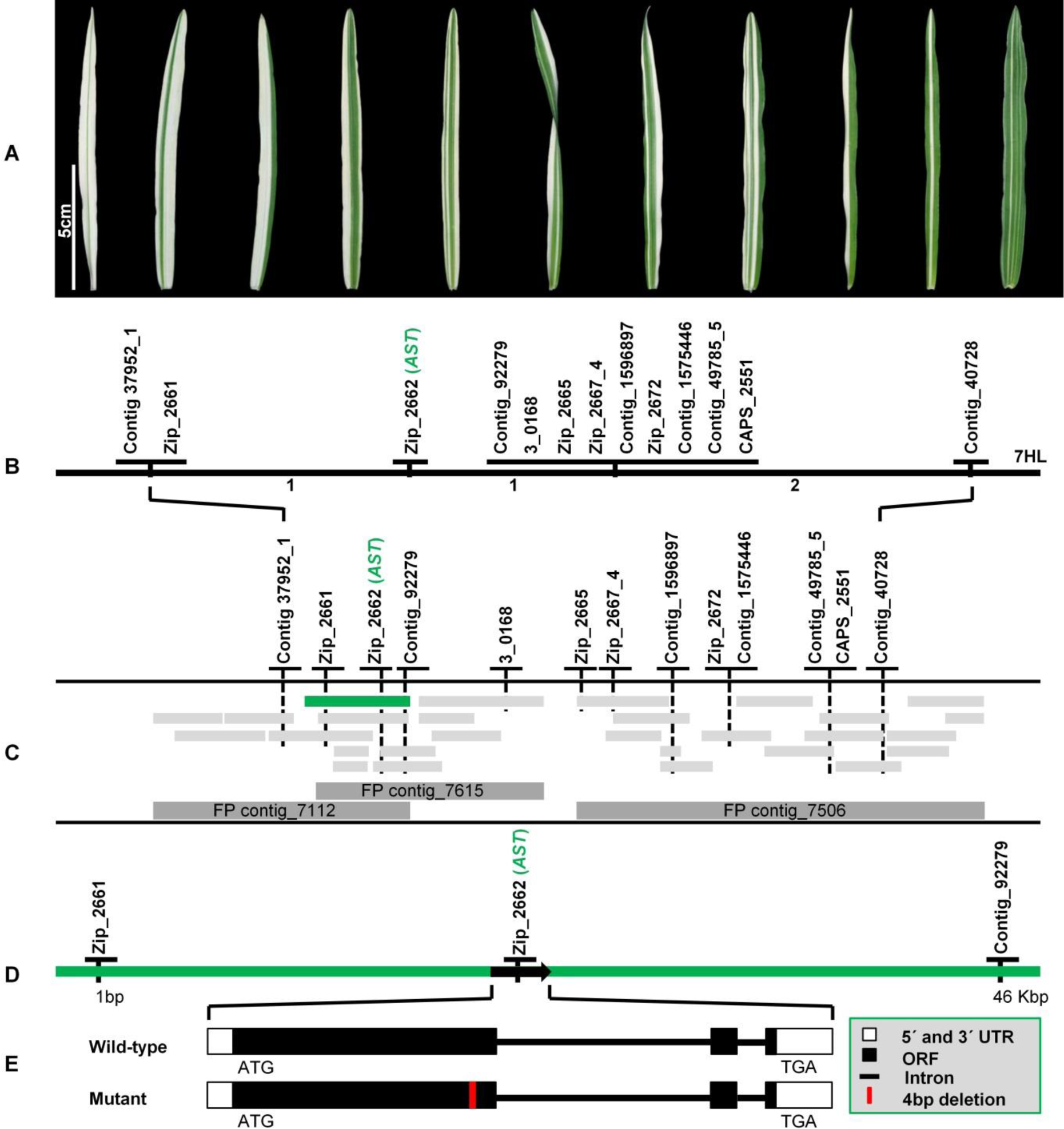
Map-based cloning of the *HvAST* gene. **(A)** Examples of variegation of leaf coloration of the homozygous *albostrians* mutant plants. **(B)** High-resolution genetic mapping of the *HvAST* gene. One co-segregating marker (Zip_2662) was identified and the recombination between contiguous markers was indicated by the numbers below. **(C)** Physical anchoring of markers to the sequenced MTP BACs. The two flanking markers Zip_2661 and Contig_92279 as well as the co-segregating marker Zip_2662 were located on the same BAC clone HVVMRXALLrA0395M21 (FP contig_7112) as indicated in green color. Each grey bar represents one BAC clone. FP contig: Finger printed contig, defined as a set of overlapping BAC clones. **(D)** Gene prediction based on repeat masked BAC assemblies of the target interval through alignment to barley gene models published by IBSC (2012). A single gene, *MLOC_670* (GenBank accession ID: AK366098) as indicated by the arrow, was identified within the target interval between two flanking markers Zip_2661 and Contig_92279, flanking a physical distance of around 46 Kbp. **(E)** Structure of the gene *HvAST*: The mutant allele harbors a 4 bp deletion (red bar) near the end of the first exon as compared to the wild-type allele found in genotype Morex. The gene structure of *HvAST* is on according to cDNA analysis of barley cultivar Morex.

### Functional Validation of the *HvAST* Gene by TILLING and Test of Allelism

To verify the biological function of the gene *HvAST*, we screened an EMS induced TILLING population (cv. Barke) (Gottwald et al., 2009) for independent mutated alleles. Forty-two EMS-induced mutations, including 20 synonymous and 20 non-synonymous mutations, one 9 bp deletion and one mutation leading to a premature stop codon, were identified. Whereas no other mutant family exhibited any chlorophyll / photosynthesis related phenotype, the M_3_ progeny of the M_2_-TILLING family 6460-1 (carrying the premature stop codon) was segregating for albinism. All homozygous mutant progeny of TILLING family 6460-1 showed a complete albino phenotype (Figures 2A and 2B), while homozygous wild-type and heterozygous plants from the segregating M_3_ all represented entirely green seedlings. The linkage between albino phenotype and mutant genotype of the *HvAST* candidate gene was further validated through analysis of a large M_4_ population comprising 245 individuals derived from five M_3_ heterozygous plants (Figure 2A). All homozygous M_4_ mutants grew into purely albino seedlings. The Mendelian segregation pattern (χ^2^=0.74; *df*=2; *p*=0.69) of the M_4_ generation indicated that, like the *albostrians* phenotype in M4205, the albino phenotype was controlled by one single recessive gene. We observed exceptions from the pure albino phenotype in homozygous mutant M_4_ and M_5_ progenies of TILLING family 6460-1 in the case of two seedlings showing very narrow green sectors on the first leaf (Figures 2C) indicating an analogous, yet much more severe phenotype compared to the variegation conferred by the original *HvAST* mutation (*i.e.* the 4bp deletion). Due to the severity and predominance of albino sectors, both M_4_ and M_5_ striped plants did not develop beyond the seedling stage (Figure 2D). The pre-stop allele in the identified TILLING family 6460-1 is designated as *Hvast2*.

**Figure 2.**
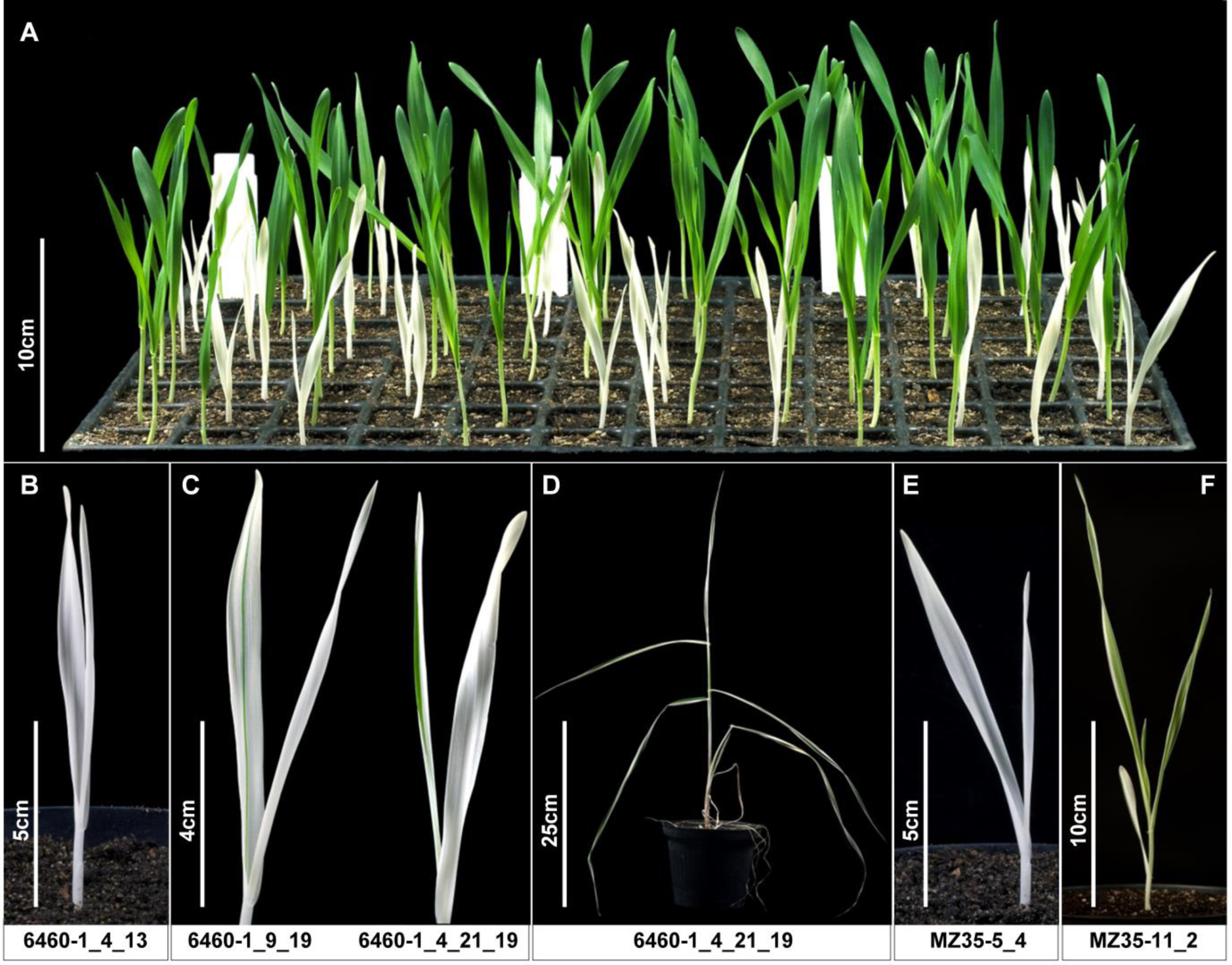
Functional validation of *HvAST* by TILLING and allelism test. **(A)** Phenotypic segregation population in M_4_ of TILLING family 6460-1. The homozygous pre-stop TILLING mutants show an albino phenotype and are delayed in growth compared with the wild-type. The plants shown here are derived from three heterozygous M_3_ plants (*i.e.* 6460-1_4, 6460-1_9, and 6460-1_11). **(B)** Example of homozygous *HvAST* pre-stop TILLING mutant, which exhibits an albino phenotype. **(C)** Examples of striped phenotype of the TILLING mutants. A single striped mutant was identified in both M_4_ generation (6460-1_9_19) and M_5_ generation (6460- 1_4_21_19). **(D)** Phenotype of the striped M_5_ plant at later developmental stage. Both variegated mutants did not reach the reproductive stage. **(E)** and **(F)** F_1_ hybrids of original *albostrians* mutant and pre-stop TILLING mutant show either albino (MZ35-5_4) or green-white striped phenotype (MZ35-11_2).

The prominent characteristic of the *albostrians* mutant is the lack of 70S ribosomes in the plastids of albinotic leaf sectors (Hess et al., 1993). We checked therefore whether in albino leaves of the allelic TILLING mutant 6460-1 plastid ribosomes are also missing. Indeed, as in *albostrians*, ribosomes could not be detected by electron microscopy in defective plastids of the TILLING mutant (Figure 3A); and the plastid rRNAs were missing in preparations from albino leaves as determined by formaldehyde agarose gel electrophoresis and evaluation using an Agilent 2100 Bioanalyzer (Figure 3B). Consequently, the development of plastids in the albino sectors of the TILLING mutant is extremely impeded, comparable in extent to *albostrians* defective plastids. In contrast to the well-developed crescent-shaped chloroplasts in the wild-type and green TILLING mutant, smaller and irregularly shaped plastids are observed in the *albostrians* and albino TILLING mutants. Instead of typical thylakoids and granum stacks as seen in wild-type chloroplasts, only a few vesicle-like structures are found within mutant plastids (Figure 3A).

**Figure 3.**
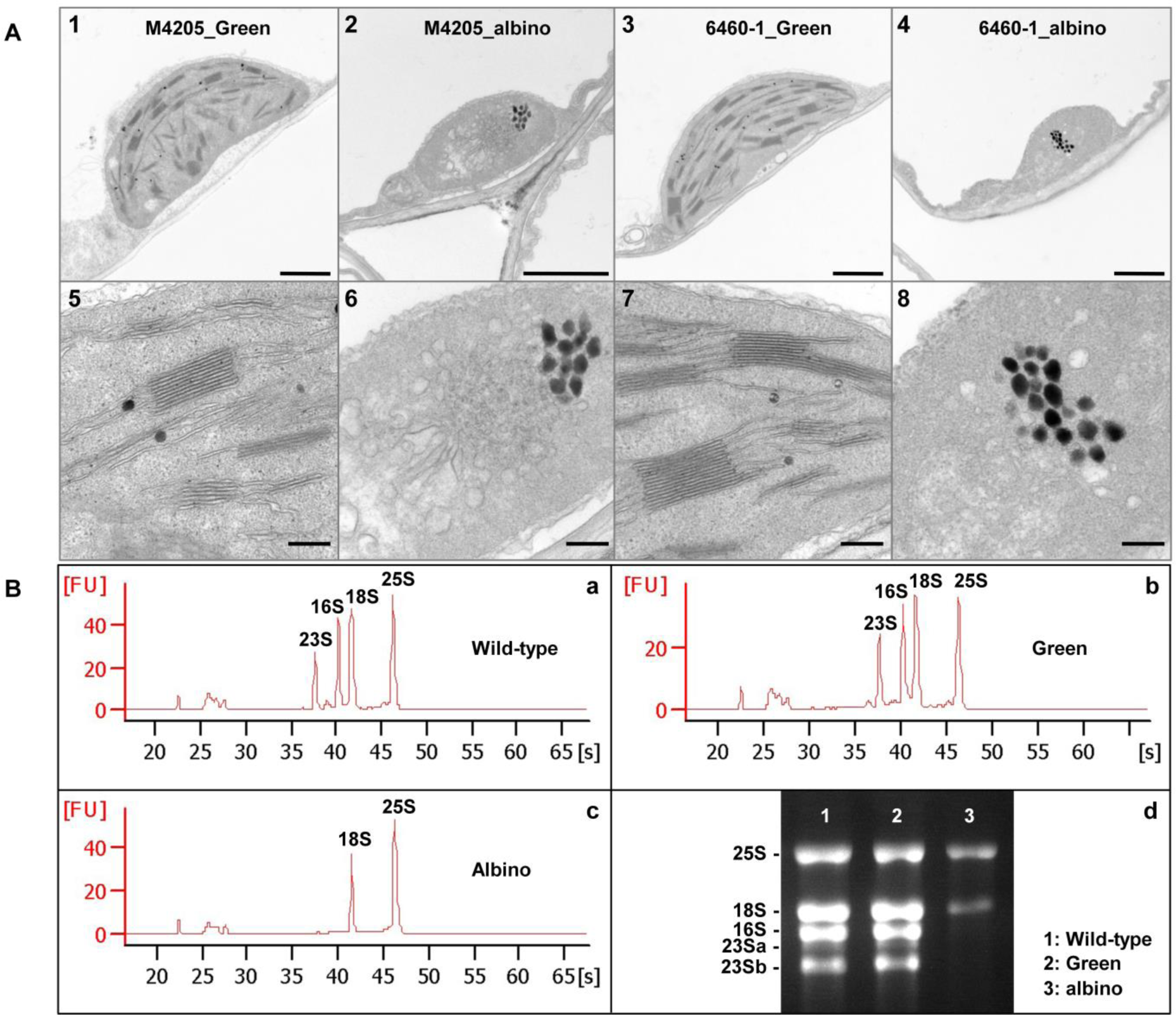
Ultrastructural analysis of the *albostrians* and TILLING mutants and analysis of rRNA from wild-type and mutant of TILLNG family 6460-1 using an Agilent 2100 Bioanalyzer. **(A)** Ultrastructural analysis of the *albostrians* and TILLING mutants. Green leaves of the *albostrians* mutant (panels 1 and 5) and wild-type (for *HvAST* gene locus) TILLING plant (panels 3 and 7) contain chloroplasts with fully differentiated grana and stroma thylakoids and ribosomes. In contrast, only some vesicle-like structures and plastoglobuli are observed in plastids of albino leaves of the *albostrians* (panels 2 and 6) and TILLING (panels 4 and 8) mutants. The lower panels represent larger magnification of the corresponding plastid in the upper panel. Scale bar for panels 1- 4: 1 μm, and panels 5-8: 200 nm. **(B)** Analysis of rRNA from wild-type and mutant of TILLNG family 6460-1 using an Agilent 2100 Bioanalyzer. Homozygous wild-type (a), green sectors of the homozygous TILLING mutant (b) and white sectors of the homozygous TILLING mutant (c). Separation of total RNA extracted from the samples mentioned above in formaldehyde agarose gel (d).

Crosses between plants carrying the original 4 bp deletion *HvAST* allele of genotype M4205 (*ast1*/*ast1*) and the novel TILLING pre-stop allele (*AST*/*ast2*) demonstrated the allelic state of both mutations. Seven of the 18 F_1_ plants showed a green phenotype as expected for plants heterozygous for the original *HvAST* allele (*AST*/*ast1*). The remaining eleven plants (*ast1*/*ast2*), heterozygous for the two mutated alleles, were either completely albino or green-white variegated (Figures 2E and 2F). This analysis confirmed that the identified candidate gene at the *MLOC_670* locus is the functional gene underlying the mutant phenotype of the original *albostrians* mutant genotype M4205. We named the gene *HvAST* [for *Hordeum vulgare ALBOSTRIANS* according to (Hagemann and Scholz, 1962; Franckowiak et al., 1992)].

### Site-directed Mutagenesis of *HvAST* by RNA-guided Cas9 Endonuclease

*HvAST* as the causal gene for the striped/albino phenotype was confirmed by the allelic nature of the *albostrians* and TILLING mutants. Remarkably, the phenotype of the homozygous TILLING mutant is much more severe than in the original *albostrians* mutant, which is probably due to the different relative location of the two mutations within the coding region (Figure 4). In an attempt to verify this presumption and to reproduce the *albostrians* phenotype, we employed site-directed mutagenesis by RNA-guided Cas9 endonuclease to generate new *HvAST* mutants close to the original site of the *albostrians* mutation of M4205. Two *HvAST*-specific guide RNAs (gRNAs) surrounding the 4 bp deletion region of the original *albostrians* mutant were selected (Figures 5B and 5C). Twenty-one out of twenty-three T_0_ regenerated plantlets carried the intact T-DNA, *i.e.* an *OsU3* promoter-driven guide RNA and a maize codon-optimized *Cas9* controlled by the maize *UBIQUITIN1* promoter with first intron. Twenty of these showed fully green leaves, whereas one plant (BG684E11, carrying the gRNA specific for target motif 1 which resides a few base-pairs upstream of the original *albostrians* mutation site) exhibited a green-white-striped phenotype resembling that of mutant M4205. A variegated leaf of BG684E11 was used for genotyping the pre-selected *HvAST*-region which revealed two mutant alleles each carrying a single base-pair (A/T) insertion within target motif 1 at the position expected to be cut by the gRNA/Cas9 complex (Figure 5D). Genotyping revealed the chimeric nature of the original T_0_ plant BG684E11, since four genotypic classes could be observed in the corresponding T_1_ progeny: homozygous *mutant 1*/*mutant 1* or *mutant 2/mutant 2* (class I), heterozygous *mutant 1/mutant 2* (class II), *mutant 1* or *mutant 2/wild-type* (class III) and homozygous *wild-type/wild-type* (class IV) (Figure 5E). The individuals of class I and II either showed green-white variegated (Figure 5A) or albino leaves, while plants in the remaining two classes exclusively were completely green. Hence, site-directed mutagenesis of *HvAST* by RNA-guided Cas9 endonuclease reproduced a phenotype similar to *albostrians*, which unambiguously confirmed *HvAST* as the functional associated gene. Our findings supported also the hypothesis that the severity of the *HvAST* mutant phenotype correlates with the relative position of lesions within the gene. The Cas9-induced mutant allele with 1 bp insertion (nucleotide G) was designated as *Hvast3*.

**Figure 4.**
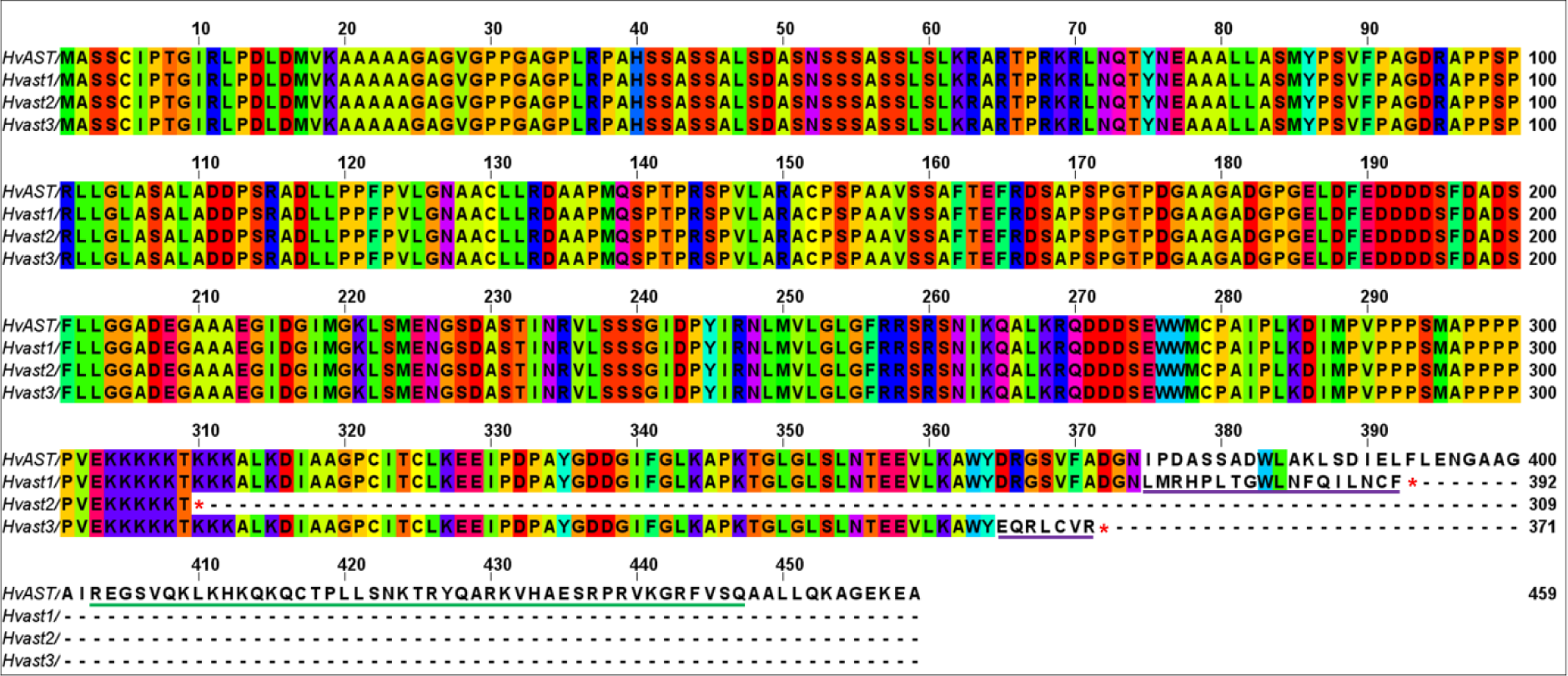
Alignment of HvAST protein sequences encoded by wild-type and mutant alleles. The position of the stop codon of the truncated proteins is indicated with a red asterisk. The altered amino acids of the truncated proteins are indicated by purple underlines. The CCT domain of HvAST protein is indicated by green underline. Alignment was performed by help of the online tool Clustal Omega and all gaps were deleted. The figure was generated by visualization under the online Jalview applet.

**Figure 5.**
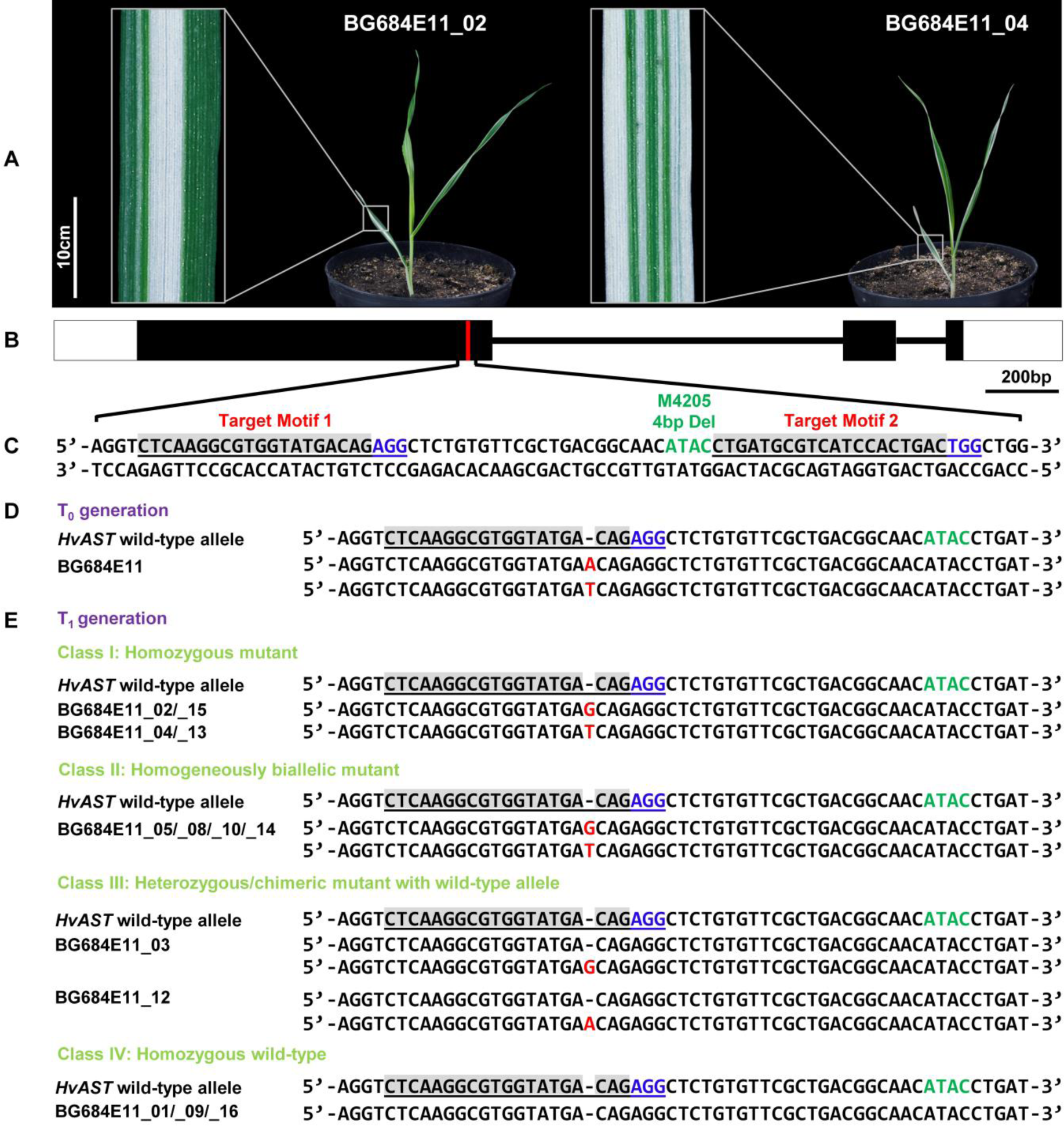
Site-directed mutagenesis of *HvAST* gene by RNA-guided Cas9 endonuclease. **(A)** Examples of the variegated homozygous *HvAST* mutants. **(B)** Gene structure of the *HvAST* gene. The region of 4 bp deletion carried by the original *albostrians* mutant is marked as red bar. **(C)** The genomic target motifs 1 and 2 (underlined) were selected up- and downstream of the 4bp deleted (indicated by green letters) in the original *albostrians* mutant. Guide RNAs were designed to address the sequences indicated by grey background. The protospacer adjacent motifs (PAM) bound by the Cas9 protein are marked with blue color. **(D)** Mutation detection in T_0_ plants. One biallelic mutant was identified which carries a one-nucleotide insertion 3 bp upstream of the PAM of target motif 1. **(E)** Inheritance of mutations in the T_1_ generation. The progenies of the T_0_ homogeneously biallelic mutant segregated into four classes of plants based on the genotypic status of the *HvAST* gene. All the homozygous (Class I) and biallelic mutants (Class II) exhibit either green-white striped or albino phenotypes, while heterozygous/chimeric mutants with wild-type allele (Class III) and homozygous wild-type plants show green phenotype.

### *HvAST* is a Member of the *CCT MOTIF FAMILY* of Genes and is Targeted to the Plastids

Sequence comparison revealed that *HvAST* encodes a putative protein with a length of 459 amino acids (AA). It carries a CCT domain at AA position 403 to 447 according to survey on the NCBI’s Conserved Domain Database (Marchler-Bauer et al., 2017). CCT is a conserved domain of 43 AA found in the *Arabidopsis thaliana* transcription factors <u>C</u>ONSTANS (CO), <u>C</u>O-LIKE, and <u>T</u>IMING OF CHLOROPHYLL A/B BINDING PROTEIN1 (TOC1) (Putterill et al., 1995; Strayer et al., 2000). The gene *HvAST* was previously described as *HvCMF7* in a study on the evolution of the CCT domain containing gene family (*CMF*) in *Poaceae* (Cockram et al., 2012). Based on public gene expression data derived from eight tissues/growth stages of barley International Barley Genome Sequencing Consortium (2012) the gene *HvAST* is expressed in all eight tissues/stages reaching the highest level of expression in young barley tillers and the lowest expression in the embryo and developing grains, respectively (Supplemental Figure 1). So far investigated, CMF proteins were proposed to represent transcription factors (Ben-Naim et al., 2006) and the CCT domain was predicted to be a nucleus localization signal (Robert et al., 1998). For HvAST, *in silico* prediction (Emanuelsson et al., 1999; Emanuelsson et al., 2000; Small et al., 2004) indicated at high probability the presence of an N-terminal chloroplast transit peptide (cTP) suggesting chloroplast targeting of the protein. We tested the subcellular localization after fusing green fluorescent protein (GFP) to the C-terminus of HvAST.

Four constructs were made for *HvAST*: wild-type *HvAST* (WT_HvAST:GFP), two mutated forms [*i.e.* the original *albostrians* mutant M4205 (M4205_HvAST:GFP) and the TILLING mutant 6460-1 (TILLING_HvAST:GFP)] and the cTP domain of HvAST (cTP_83AA_HvAST:GFP) (Figure 6A). These fusion constructs, together with one plastid marker pt-rk-CD3-999 (Nelson et al., 2007), respectively, were subjected to transient expression experiments using biolistic bombardment of barley leaf segments with vector-coated gold particles. The plastid marker pt-rk-CD3-999 (Nelson et al., 2007) was detected by its orange fluorescence (Figure 6, mCherry column). Cells expressing only the GFP (non-fusion GFP control) showed green fluorescence associated with the nucleus and the cytoplasm but not with plastids (Figure 6B). In contrast, the mCherry signal of the plastid marker accumulated exclusively in plastids (Figure 6C). WT_HvAST:GFP and its two truncated mutant forms were detected as green fluorescence in leaf epidermal cells associated with the nuclei and stronger with the plastids (Figures 6D, 6E and 6F). The plastid localization of HvAST was further confirmed by the cTP_83AA_HvAST:GFP, which showed strong accumulation of GFP signals in the plastids (Figure 6G). In total, 30-50 cells for each of the five fusion constructs were checked for the presence of both green and orange fluorescence.

**Figure 6.**
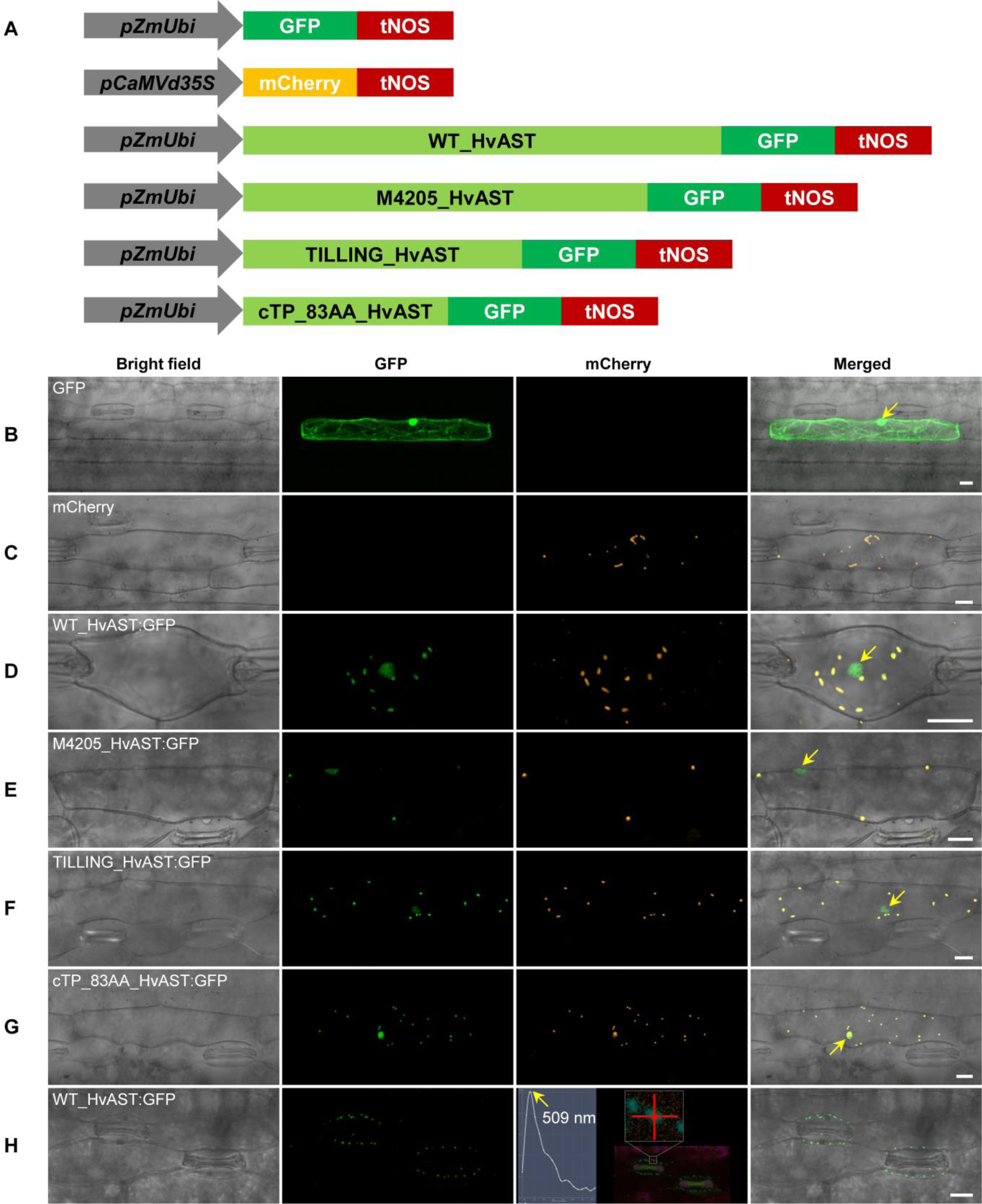
Subcellular localization of HvAST:GFP fusion proteins. **(A)** Schematic drawing of the constructs used in biolistic bombardment. *pZm*Ubi: maize *UBIQUITIN1* promoter. *pCaMV*d35S: Cauliflower Mosaic Virus double *35S* promoter. mCherry: mCherry fluorescent protein. GFP: green fluorescent protein. tNOS: *Agrobacterium NOPALINE SYNTHASE* terminator. WT_HvAST: Coding sequence of wild-type *HvAST* gene. M4205_HvAST: Coding sequence of mutant allele *Hvast1* of the original *albostrians* mutant line M4205. TILLING_HvAST: Coding sequence of mutant allele *Hvast2* of the pre-stop TILLING mutant line 6460-1. cTP_83AA_HvAST: N-terminal chloroplast transit peptide of HvAST with a length of 83 amino acids as predicted by online tool PredSL (Petsalaki et al., 2006). The schematic drawing is not in proportion with gene length. **(B)** Bombardment using gold particles coated with pSB179 plasmid (expressing GFP driven by the maize *UBIQUITIN1* promoter) as control. The green fluorescence is observed in the nucleus and the cytoskeleton / cytoplasm. **(C)** Biolistic assay for plastid marker pt-rk-CD3-999 which carries the *mCherry* gene driven by a double *35S* promoter. The mCherry signal (orange color) exclusively accumulates in the plastids. **(D)** Particle co-bombardment of both plastid marker pt-rk-CD3-999 and wild-type fusion construct WT_HvAST:GFP. The bright field, GFP, and mCherry signals are displayed as individual channels and the merged channels are shown on the rightmost panel. GFP and mCherry fluoresce green and orange, respectively. The green fluorescence of the wild-type fusion protein accumulates in plastids and nucleus of the barley epidermal cells. Thus, HvAST protein targeted to plastid; the nucleus signal, nevertheless, cannot be resolved since GFP alone also shows clear strong accumulation in the nucleus. **(E)** and **(F)** Subcellular localization of the two mutant proteins M4205_HvAST:GFP **(E)** and TILLING_HvAST:GFP **(F)** was also investigated by co-bombardment with the plastid marker pt-rk-CD3-999 and display the same subcellular localization as the wild-type protein **(D)**. **(G)** The chloroplast localization of HvAST was further confirmed by the fact that its N-terminal (83 AA chloroplast transit peptide) drives the GFP protein into plastids. **(H)** Subcellular localization of WT_HvAST:GFP in barley stable transgenic lines. Consistent with the results from transient expression, the GFP fluorescence, characterized by photospectrometic analysis with an emission peak at wavelength 509 nm, was specifically accumulated in the plastids. The yellow arrows in the merged panels indicate the nucleus. The first leaf of 10- day-old barley seedlings was used for particle bombardment. The fluorescence was checked 24 hours after bombardment. Scale bar for all images is 20 μm.

To further confirm subcellular localization of HvAST, stably transgenic barley lines carrying WT_HvAST:GFP were obtained through *Agrobacterium*-mediated transformation. Consistently with results of the transient experiments, GFP fluorescence was specifically targeted to the plastids in the epidermal cells (Figure 6H). Surprisingly, we could not detect GFP fluorescence in the mesophyll cells.

Taken together, the wild-type HvAST and its two mutated forms displayed a clear compartmentalized accumulation in the plastids. Although also nuclear fluorescence was visible in the transformed cells, which would support a role of the CCT domain as a nuclear localization signal, further investigations are required to discriminate if the observed localization of HvAST to the nucleus is only the consequence of unspecific nuclear targeting of GFP as was reported in other systems before (Seibel et al., 2007).

## DISCUSSION

Leaf variegation in the barley mutant *albostrians* is distinct due to the incomplete penetrance of the phenotype. Even if carrying the mutant allele at homozygous state, plants may be completely green and the progeny of such a green plant will segregate in a ratio of 1:8:1 for green, variegated, or albino phenotype, respectively. This phenomenon implies residual activity of the affected allele and a threshold mechanism involved at a critical decision point in chloroplast development / maturation - a hypothesis, which can be further tested now, since the gene *HvAST* was isolated by positional cloning.

The cloning of the gene *HvAST* was supported by four lines of evidence: (i) the smallest genetic interval contained only a single gene in perfect linkage with the phenotype, (ii) mutant analysis by TILLING revealed an independent pre-stop allele leading to a related but more severe and fully penetrant phenotype, (iii) crossing of the two independent mutant alleles proved their state of allelism, and last not least (iv) *albostrians*-like alleles were induced by site-directed mutagenesis using RNA-guided Cas9 endonuclease. In the original *albostrians* mutant allele, *ast1* of the genotype M4205, a four base pair deletion at position 1123-1126 of the coding sequence is predicted to induce a shift of reading frame and, as a consequence, a premature stop codon in the second exon of the gene (Figure 4). During functional validation of the identified gene, the characterization of two novel independent mutant alleles, *ast2* and *ast3*, implied that severity and penetrance of the phenotype is correlated with the relative position of the mutation as well as with allele dosage. The albino phenotype conferred by *ast2*, induced by EMS and identified by TILLING, suggested (almost) complete loss of protein activity. Though this phenotype is not completely identical with the mostly striped progeny of *albostrians* barley, *ast2* provided evidence for the identified gene *HvAST* representing *albostrians*, since also two albino seedlings with tiny green stripes could be observed among 289 M_3_, M_4_ and M_5_ progenies. F_1_ plants heterozygous for *ast1*/*ast2* confirmed their allelic state but also demonstrated that the phenotype is affected by allele dosage. In contrast to M4205 (*ast1*/*ast1*), F_1_ plants (*ast1*/*ast2*) were either albino or green-white variegated but not a single fully green F_1_ could be observed. Since plants heterozygous with one wild-type allele (*AST*/*ast1*) are generally fully green, the severity of phenotype thus varies in an allele-dosage dependent manner. The phenotypic difference of *ast1* and *ast2* homozygous mutants provided the first indication of a correlation between relative position of the mutation and severity of the phenotype. This hypothesis was confirmed by a third allele, *ast3*, obtained via site-directed mutagenesis using RNA-guided Cas9 endonuclease. A one-nucleotide insertion at 28 bp upstream of the 4 bp deletion in *ast1* led to a frameshift and an *ast1-*similar gene product of seven amino acids longer altered putative C-terminal sequence. Homozygous seedlings carrying this mutation (*ast3*/*ast3*) resembled the typical *albostrians* green-white striped or albino seedling phenotype, however, in contrast to *ast1*/*ast1* plants, no green seedlings were observed indicating a more severe phenotype for *ast3*/*ast3* plants.

### HvAST is a CCT-domain Protein and Homolog of the Arabidopsis CIA2 Protein

*In silico* analyses of the amino acid sequence of HvAST revealed at its N-terminus a putative chloroplast transit peptide and near the C-terminus a CCT-domain qualifying it to be a member (HvCMF7) of the larger CCT domain-containing gene family of the *Poaceae* (Cockram et al., 2012). The CCT-domain is named after rather well studied plant transcription factors: CONSTANS, CONSTANS-LIKE, and TOC1 (TIMING OF CHLOROPHYLL A/B BINDING PROTEIN1) (Strayer et al., 2000). *HvAST* belongs to a sub-gene family that is characteristic for carrying only a single CCT domain per gene (Cockram et al., 2012) and CCT is the only domain of HvAST in common with these transcription factors.

The closest homologs of *HvAST* in *Arabidopsis thaliana* are the genes “*CHLOROPLAST IMPORT APPARATUS 2* (*AtCIA2*)” and “*CHLOROPLAST IMPORT APPARATUS 2-LIKE (AtCIL)*” (Sun et al., 2001). *AtCIA2* encodes a nucleus-localized transcription factor involved in expression of both protein translocon and ribosomal protein genes by binding to the respective promoters (Sun et al., 2009a). Yet, *HvAST* in barley might have a different function. As expected by the *in silico* predicted chloroplast transit peptide, it could be shown that HvAST was localized to barley chloroplasts in transient co-localization and stable transformation experiments. GFP signal was also observed in the nucleus, which, however, is not providing yet any evidence of dual targeting to the plastid and nucleus, since GFP has a general tendency of nuclear localization as demonstrated by the GFP-only controls in the transient expression experiments. Searching the public plant proteomics database (Sun et al., 2009b) revealed no *HvAST* homologs of other plant species targeted to the plastids. Thus, to our knowledge, HvAST represents the first CCT domain protein found to be targeted to plastids.

Nevertheless, the exact molecular function of HvAST remains elusive. Based on the analysis of independent *HvAST* mutant alleles, it can be concluded that the CCT domain may be involved in overall HvAST-function, however, this domain is not exclusively required to regulate proper chloroplast differentiation or maturation in barley. The alleles *ast1* and *ast3* allow, at least in a threshold dependent manner, the formation of fully functional chloroplasts in green seedlings and green leaf sectors, hence normal differentiation of chloroplasts despite complete loss of the CCT domain. Both mutated alleles reduce most likely the protein functionality but do not completely destroy its function. The increasingly severe phenotype in mutant alleles with defects progressing towards the N-terminus of the gene indicated the presence of an additional, still to be described domain of the protein that is involved in communicating its essential function in the different cell compartments. Further protein-protein and DNA-protein interaction studies involving the series of barley *HvAST* mutants will be required to further address this question.

It is interesting to note that we found phenotypic effects only for mutations that affect large portions of the HvAST protein at its C-terminus. The TILLING population contained also several mutations that introduced putative synonymous or non-synonymous exchanges of amino acids into the protein sequence. All of them showed the wild-type phenotype. The gradual increase in the amount of white leaves and white leaf sectors with shifting the position of the mutation toward the N-terminus of the HvAST protein and the absence of effects of point mutations leading to an altered amino acid in this region suggests that the C-terminal part of the protein does not have a catalytic function but potentially rather interacts with other proteins – potentially involved in expression or regulation of genes required for plastid ribosome formation. In case of the mutant *Hvast* alleles, this interaction would be less efficient or missing suggesting the formation of a multiprotein-complex with less or no remaining activity.

### HvAST – A Factor Involved in Chloroplast Ribosome Formation?

Remarkably, green, green-white striped and white seedlings of the *albostrians* mutant have all the same genotype; they are homozygous for the recessive mutant allele (*ast1*) and they have identical plastid genomes (Hess et al., 1993; Zhelyazkova et al., 2012). The different fate of plastids in *albostrians* leaves is therefore not caused by a mutation in the DNA of those proplastids that do not develop into chloroplasts. There is no environmental influence on the phenotype of *albostrians* plants contrasting reports on other variegated plants with identical genotype in differently colored leaf sectors like barley *tigrina* mutants or the Arabidopsis *immutans* mutant. In these mutants the pigmentation of leaves and leaf sectors depends on the light conditions, hence illumination induces chlorosis due to photo-oxidation (Wettstein et al., 1974; Wetzel et al., 1994; Lee et al., 2003; Putarjunan et al., 2013). Most certainly, it is the ribosome deficiency of all or only part of the plastids, respectively, in the meristems of the first and following leaves that blocks further chloroplast development and thus either causes the formation of completely white or green-white variegated leaves, respectively. As was demonstrated before (Knoth and Hagemann, 1977; Dorne et al., 1982) and confirmed in the present study, plastids in white leave sectors of *albostrians* plants do not contain ribosomes, thus HvAST is needed directly or indirectly for the biogenesis of plastid ribosomes. Since HvAST or its homologs in Arabidopsis have not been described as a component of plastid ribosomes, *i.e.* as a ribosomal protein *sensu stricto* (Yamaguchi and Subramanian, 2000; Yamaguchi et al., 2000; Tiller et al., 2012), the HvAST protein might not be associated with ribosomes or be associated only under certain conditions and might be required for the assembly of plastid ribosomes rather than for their function.

### Conclusion

The identification of the barley gene *HvAST* has revealed novel insights into the role of CCT-domain containing proteins. We show that a barley CCT-domain containing protein is localized to plastids. This gene likely plays a crucial role during early embryo development for plastid ribosome formation and hence for chloroplast development. An Arabidopsis homolog of *HvAST* was previously published to act as transcriptional regulator for nuclear genes coding for chloroplast ribosomal proteins and for the chloroplast protein translocon (Sun et al., 2009a). Based on our findings, a dual role of this special group of CCT domain proteins in nuclei and in plastids cannot be ruled out. The identification of the gene causing the “*albostrians*” phenotype will foster studies on early phases of chloroplast development, in particular the formation of plastid ribosomes, on possible dual localization of CCT proteins to nuclei and plastids, and on leaf variegation in non-chimeric plants.

## METHODS

### Plant Materials

The six-rowed spring barley cultivar ‘Morex’ and the two-rowed spring barley cultivar ‘Barke’ were used as maternal parent in crossings with the mutant line M4205 (Hagemann and Scholz, 1962), respectively. The obtained F_1_ were self-pollinated to obtain F_2_ populations for genetic mapping. The two F_2_ mapping populations were designated as ‘MM4205’ (‘Morex x M4205’) and ‘BM4205’ (‘Barke x M4205’) indicating the respective parental combinations of the crosses. All plants were grown under greenhouse conditions with a day/night temperature and photoperiod cycle of 20°C/15°C and 16h light / 8h darkness, respectively.

### DNA Preparation

DNA was isolated according to the protocol of Doyle and Doyle (1990) from leaf samples collected from three-leaf stage greenhouse-grown seedlings and quantified using NanoDrop Spectrophotometer (Thermo Scientific, Wilmington, USA) according to the manufacturer’s instructions and adjusted to 100 ng/µl for any PCR application.

### PCR Reaction

DNA amplification reactions were performed in a total volume of 20 µl containing 40 ng of template DNA, 4 mM of dNTPs, 1 µl each of 5 µM forward and reverse primer, 0.5 units of HotStarTaq DNA polymerase (Qiagen, Düsseldorf, Germany) and 2 µl of 10x PCR buffer (100 mM Tris-HCl, pH 8.3; 500 mM KCl; 15 mM MgCl_2_; 0.01% gelatin). Touch-down PCR program was used with a GeneAmp 9700 thermal cycler (Life Technologies GmbH, Darmstadt, Germany): initial denaturation at 94°C for 15 min followed by 5 cycles at 94°C for 30 s, annealing at 65°C to 60°C (−1°C/cycle) for 30 s, extension 1 min at 72°C, and then proceeded for 40 cycles 94°C for 30 s, 60°C for 30 s, 72°C 1 min, and followed by a final extension at 72°C for 10 min. Detailed information of the primers used for marker development is presented in Supplemental Table 1.

### CAPS Assay

SNP polymorphisms identified between the mapping parents were converted into CAPS (Cleaved Amplified Polymorphic Sequences) markers (Thiel et al., 2004) (Supplemental Table 1). For this purpose, PCR products were purified using the NucleoFast^®^ 96 PCR Kit (Macherey-Nagel, Düren, Germany) and sequenced on ABI 3730 XL (Life Technologies GmbH, Darmstadt, Germany). Sequences were aligned by using Sequencher^®^ version 5.2.3 software (Gene Codes Corporation, Ann Arbor, MI USA. http://www.genecodes.com) for SNP identification. Subsequently, SNP2CAPS software (Thiel et al., 2004) was adopted to select a suitable restriction endonuclease. The resulting fragments were resolved by electrophoresis on 1.5% (w/v) agarose, 1x TBE gels (Invitrogen GmbH, Darmstadt, Germany).

### Genetic Mapping

As an initial step, 91 F_2_ individuals from each population ‘MM4205’ and ‘BM4205’ were selected for low resolution genetic mapping the gene *HvAST*. Genotyping was performed by using the Illumina Golden Gate assay with a custom set of 381 BOPA SNP markers (Close et al., 2009). Subsequently, only the ‘MM4205’ population was further employed for fine mapping of the gene *HvAST* due to its higher rate of polymorphic markers compared to the ‘BM4205’ population. Saturation mapping within the ‘MM4205’ population (91 genotypes) was conducted in an effort to narrow down the target genetic interval. Next, fine mapping the gene *HvAST* with an additional 1344 F_2_ plants was scheduled in two steps. First, 142 recombinants were selected by screening 960 F_2_ plants with flanking markers CAPS_2536 and CAPS_2560 and used for further marker saturation. In a second step, 384 variegated or albino F_2_ individuals were selected from 1920 F_2_ plants and screened for recombination between newly identified flanking markers Zip_2661 and Zip_2680_1. In addition, the genotype at the *HvAST* locus was determined by phenotyping 30 F_3_ seedlings derived from each F_2_ recombinant through self-pollination. While wild-type F_2_ plants are expected to yield 100% green progeny, F_3_ families derived from heterozygous F_2_ will segregate into 75% wild-type and 25% variegated or albino plants. F_3_ progeny of F_2_ homozygous green mutants will segregate into 10% green, 80% variegated, and 10% albino seedlings. All molecular markers used for saturation mapping as well as for fine mapping relied on publicly available genomic resources (Sato et al., 2009; Mayer et al., 2011; International Barley Genome Sequencing Consortium, 2012; Mayer et al., 2012).

### Linkage Analysis

Genetic linkage analysis was performed using JoinMap 4 software (Van Ooijen, 2006). Homozygous wild-type, heterozygous and homozygous mutant allele calls were defined as A, H and B, respectively; missing data was indicated by a dash. Maximum Likelihood algorithm and Kosambi’s mapping function were chosen for building the linkage maps. Markers were assigned into seven groups based on Logarithm of Odds (LOD = 4) groupings. Visualization of maps derived from JoinMap 4 was achieved by MapChart software (Voorrips, 2002).

### Physical Mapping

The target physical region was identified by anchoring the flanking genetic markers to the physical map of barley (International Barley Genome Sequencing Consortium, 2012). Two anchoring strategies were applied: (1) *in silico* anchoring approach through BLASTn (Mount, 2007), *i.e.* sequence of the markers were used to screen by BLAST against the draft barley sequence assembly at IPK barley BLAST Server (http://webblast.ipk-gatersleben.de/barley/) or the HarvEST Server (http://138.23.178.42/blast/index.html). (2) Experimental anchoring via PCR screening of a BAC library derived from barley cultivar ‘Morex’ (Schulte et al., 2011) as previously described (Ariyadasa and Stein, 2012) (Supplemental Table 2). The MTP (Minimal Tiling Path) BACs spanning the region represented by the physical map contigs were shotgun sequenced (Beier et al., 2016) on the Miseq^®^ System (Illumina MiSeq^®^, San Diego, CA, USA) (Supplemental Table 3). The shotgun reads were assembled as described previously (International Barley Genome Sequencing Consortium, 2012). Gene models were predicted on non-repetitive sequences (Schmutzer et al., 2014) of the target interval through alignment of gene models defined on the Morex WGS assembly (International Barley Genome Sequencing Consortium, 2012).

### TILLING Screening

An ethylmethanesulfonate (EMS) induced TILLING population, comprising 7,979 M_2_ plants, derived from a two-rowed malting barley cultivar ‘Barke’ (Gottwald et al., 2009), was used for identification of independent mutated alleles of the gene *HvAST*. Primers were designed covering the coding sequence of the gene *HvAST* (Supplemental Table 4). PCR amplicons were analyzed combined with dsDNA Cleavage Kit (DNF-480-3000) and Gel-dsDNA reagent kit (DNF-910-K1000) according to the manufacturer’s protocols (Advanced Analytical Technologies GmbH, Heidelberg, Germany). Subsequently, the cleaved PCR products were separated using the *Advan*CE™ FS96 capillary electrophoresis system (Advanced Analytical Technologies GmbH, Heidelberg, Germany) and results were interpreted with assistance of the PRO Size™ software (Advanced Analytical Technologies GmbH, Heidelberg, Germany). The identified M_2_ TILLING mutants were confirmed by Sanger sequencing of PCR amplicons derived from the respective families. Plant families carrying non-synonymous mutations, deletions, or immature stop codons were selected for propagation. Phenotyping was performed through M_3_ to M_5_ generation of the identified M_2_ mutants. Heterozygous plants were propagated and maintained for further reproduction. All the identified TILLING families carrying mutation for the *HvAST* gene are summarized in Supplemental Table 5.

### *HvAST* Gene Structure Analysis

Primary leaf was collected from seedlings 4 days after germination. Total RNA was extracted using TRIzol^®^ reagent (Invitrogen GmbH, Darmstadt, Germany) following manufacturer’s instructions. The concentration of the obtained RNA was determined by help of a Qubit^®^ 2.0 Fluorometer (Life Technologies GmbH, Darmstadt, Germany) according to manufacturer’s manual. Genomic DNA was removed from RNA preparations by incubation with RNase-free DNase I (Fermentas, St. Leon-Rot, Germany) following the manufacturer’s instructions. The reactions were carried out in a total volume of 10 μl containing 1 μg of RNA, 1 μl of 10x reaction buffer with MgCl_2_ (100 mM Tris-HCl, pH=7.5; 25 mM MgCl_2_; 1 mM CaCl_2_) and 1 unit of DNase I (1 U/μl). Samples were incubated at 37°C for 30 min, followed by adding 1 μl of 50 mM EDTA and further incubation for 10 min at 65°C. The DNase I treated RNA was then used as template for cDNA synthesis. Reverse transcription was performed using the SuperScript™ III First-Strand Synthesis SuperMix for qRT-PCR (Invitrogen GmbH, Darmstadt, Germany) following the manufacturer’s protocol. The DNase-I treated RNA was used as template in a total volume of 30 μl containing 15 μl of 2x RT reaction mix, 3 μl of RT enzyme and 2 μl of DEPC-treated water. Reverse transcription reaction were performed during the following cycling profile: 25°C for 10 min, 50°C for 30 min, 85°C for 5 min and finally hold at 4°C. Subsequently, the RNA strand was removed from obtained cDNA by incubating at 37°C for 20 min after adding 1 μl of *E.coli* RNase H. RT-PCR was set up in a total volume of 20 μl containing 2 μl of cDNA template, 2 μl of 10x PCR buffer (100 mM Tris-HCl, pH 8.3; 500 mM KCl; 15 mM MgCl_2_; 0.01% gelatin), 2 μl of dNTPs (40 mM), 1 μl each of 5 μM forward and reverse primer, 0.1 μl of HotStarTaq DNA polymerase (5 U/μl; Qiagen, Düsseldorf, Germany) and 11.9 μl of nuclease-free water. After purification, PCR product was sequenced by using Big Dye Terminator chemistry and an ABI 3730 XL instrument (Life Technologies GmbH, Darmstadt, Germany). The *HvAST* gene structure was resolved by aligning the sequenced *HvAST* coding sequence to genomic sequence of barley cv. Morex.

### Site-directed Mutagenesis by RNA-guided Cas9 Endonuclease

The coding region of the gene *HvAST* of barley cultivar ‘Golden Promise’ was sequenced and used for gRNA/Cas9 target motif selection and guideRNA design. The search for genomic target motifs was focused to the region surrounding the 4 bp deletion carried by the original *albostrians* mutant and one proper target on each side of the deletion was selected through *in silico* analysis using the online prediction tool (https://www.deskgen.com/guidebook/; the ‘KNOCKIN’ panel was chosen for gRNA design) (Doench et al., 2014). A synthetic double-stranded oligonucleotide carrying the target-specific part of the gRNA was inserted between the *Os*U3 (RNA polymerase III) promoter and the downstream gRNA scaffold present in the monocot-compatible intermediate vector pSH91 (Budhagatapalli et al., 2016). Next, the fragment containing the expression cassettes of gRNA and Cas9 was introduced into the binary vector p6i-d35S-TE9 (DNA-Cloning-Service, Hamburg, Germany) using the *Sfi*I restriction sites. The vectors constructed for the two target motifs were then used in *Agrobacterium*-mediated co-transformation of barley cultivar ‘Golden Promise’ for generation of primary mutant plants. Presence/absence of T-DNA of the regenerated plantlets and mutation detection was achieved by PCR using custom designed T-DNA-specific and *HvAST*-specific primers, respectively (Supplemental Table 4).

### Formaldehyde Agarose Gel Electrophoresis

Electrophoresis was performed under RNase-free conditions - the electrophoresis chamber and comb were washed with 0.1% (v/v) DEPC-H_2_O and the agarose gel, which contained 2% (w/v) agarose, 1X 3-(N-morpholino) propanesulfonic acid (MOPS) buffer and 6.29% (v/v) formaldehyde, was prepared with 0.1% (v/v) DEPC-H_2_O. The RNA sample (1-5 μg) was mixed with formaldehyde loading dye, which contained 25 μl formamide (Carl Roth GmbH, Karlsruhe, Germany), 5 μl 10x MOPS buffer (200 mM MOPS; 50 mM Sodium acetate; 10 mM EDTA; Carl Roth GmbH, Karlsruhe, Germany) and 10 μl 37% formaldehyde (Carl Roth GmbH, Karlsruhe, Germany), and incubated 5 min at 65°C for denaturation, followed by adding 2 μl ethidium bromide (10 mg/ml; Carl Roth GmbH, Karlsruhe, Germany) to the mix and electrophoresis in 1x MOPS buffer at 85 V for 2.5 hours. RNA was visualized in the gel by excitation under UV light using the BioDocAnalyze Gel-analyze System (Biometra GmbH, Göttingen, Germany).

### Transmission Electron Microscope Analysis

The first leaf of seedlings at 3 days after germination was collected and three independent plants were sampled for each genotype representing three biological replicates. Leaf section preparation, fixation and embedding followed the protocol as described elsewhere with minor modification (Schwarz et al., 2015). Instead of using the Lowicryl HM20 resin, Spurr resin (Sigma-Aldrich Chemistry GmbH, Munich, Germany) was used for embedding. Ultrathin leaf sections of approximately 70 nm were used as probe for ultrastructural analysis by help of the transmission electron microscope FEI Tecnai G^2^-Sphera 200 KV (Thermo Fisher Scientific, Oregon, USA).

### Subcellular Localization

Either of the cTP domain of *HvAST*, the coding sequence of *HvAST* from wild-type (cv. Haisa), *albostrians* mutant M4205 (cv. Haisa) or TILLING mutant 6460-1 (cv. Barke) were fused to the N-terminus of the GFP reporter gene (Chiu et al., 1996) through ligation into the *Spe*I/*Hind*III cloning sites of the vector pSB179 (provided by the Kumlehn lab), respectively. To verify the potential chloroplast localization of HvAST, one plastid marker pt-rk-CD3-999 which fusions of the targeting sequence (first 79 AA) of the small subunit of tobacco rubisco to gene of mCherry fluorescent protein was employed, the expression cassette is driven by a Cauliflower Mosaic Virus 35S promoter with dual enhancer elements (d35S). Detailed information of the plastid marker can be found under https://www.arabidopsis.org/servlets/TairObject?type=stock&id=3001623338. The alternative forms of HvAST:GFP plasmids were tested via particle bombardment using PDS-1000/He system with Hepta™ adaptor Particle Delivery System (Bio-Rad Laboratories GmbH, Munich, Germany). The physical parameters, 1100 psi helium pressure and 27 inch Hg vacuum had been set to reach efficient bombardment conditions for barley epidermal cells. Preparation and delivery of DNA-coated gold particles was performed according to (Budhagatapalli et al., 2016). The sample was then scanned for the presence of fluorescence signals by help of a Confocal Laser Scanning Microscopy LSM 780 (Carl Zeiss MicroImaging GmbH, Jena, Germany).

### *Agrobacterium*-mediated Barley Transformation

Expression cassette of intermediate vector WT_HvAST:GFP, used for bombardment experiment, was cloned into binary vector p6i-d35S-TE9 (DNA-Cloning-Service, Hamburg, Germany) using the *Sfi*I restriction sites. The derived vector was designated as pML14. Subsequently, plasmid pML14 was used in *Agrobacterium*- mediated transformation of barley cultivar ‘Golden Promise’ following the protocol described elsewhere (Hensel et al., 2009).

### Accession Numbers

Sequence of the *HvAST* gene is submitted to the European Nucleotide Archive with accession number PRJEB22029. Accession number of BAC assembly for each sequenced BAC clone is summarized in Supplemental Table 3.

### Supplemental Data

**Supplemental Table 1.** Markers used for genetic mapping.

**Supplemental Table 2.** Anchoring markers to the physical map of barley.

**Supplemental Table 3.** List of the sequenced MTP BACs.

**Supplemental Table 4.** Primers used in this study.

**Supplemental Table 5.** Identified TILLING mutants for *HvAST*.

## ACKNOWLEDGMENTS

We gratefully acknowledge M. Ziems, J. Pohl, S. König and I. Walde (IPK) for their technical support in keeping plant material, performing TILLING analyses, and Sanger sequencing; H. Mueller (IPK) for photography; S. Sommerfeld (IPK) for barley transformation; M. Benecke and K. Hoffie for their help in electron microscopy analysis; K. Lenz and N. Mehlitz (HUB and IPK) for initial mapping experiments; H. Trautwein (IPK) for his help in particle bombardment experiments; A. Graner, R. Zhou, M. Jost, S. Hiekel and N. Wendler (IPK) for helpful discussions. The work was supported by a fellowship of the China Scholarship Council to M. Li, by Humboldt-Innovation GmbH (SK043) to T. Börner, by grants STE 1102/13-1 and KU 1252/8-1 of the German Research Foundation (DFG) to N. Stein and J. Kumlehn, respectively, and core funding of the Leibniz Institute of Plant Genetics and Crop Plant Research (IPK).

## AUTHOR CONTRIBUTIONS

N.S., T.B. and J.K. designed research; M.L., G.H., M. Melzer, N.B, T.R., A.H. and V.K. performed experiments; M.L., M. Mascher, and S.B. analyzed data; and M.L., T.B. and N.S. wrote the paper.

