## Supplemental file for "Leaf Variegation and Impaired Chloroplast Development Caused by a Truncated CCT Domain gene in *albostrians* Barley"

**Supplemental Data**

**Supplemental Figure 1.** Expression profile of *HvAST* in barley.
**Supplemental Table 1.** Markers used for genetic mapping.

**Supplemental Table 2.** Anchoring markers to the physical map of barley.

**Supplemental Table 3.** List of the sequenced MTP BACs.

**Supplemental Table 4.** Primers used in this study.

**Supplemental Table 5.** Identified TILLING mutants for *HvAST*.

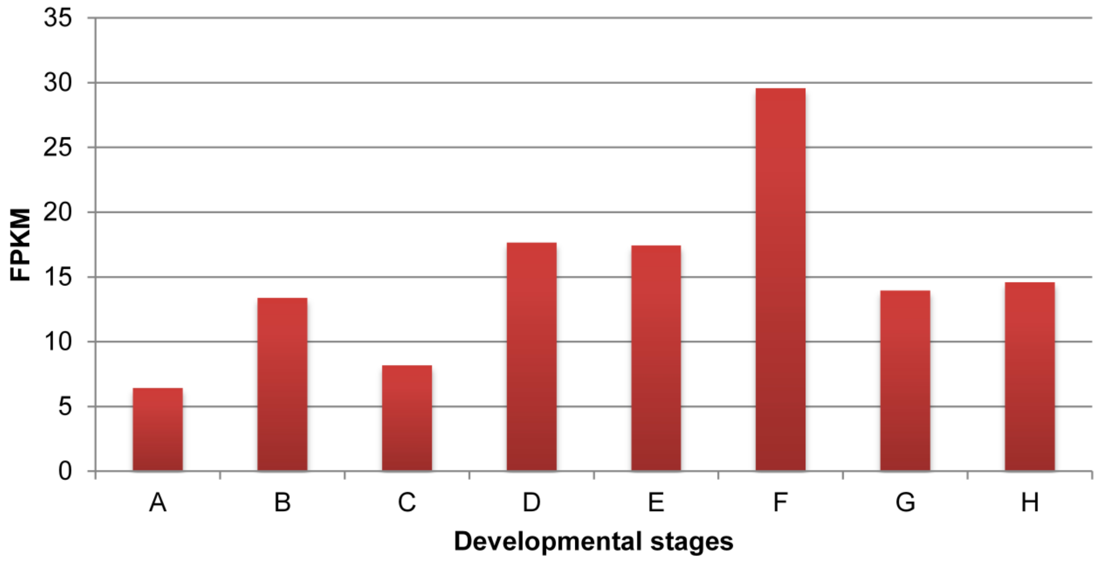


**Supplemental Figure 1:** Expression profile of *HvAST* in barley. The expression level was given as fragments per kilobase of exon per million reads mapped (FPKM) across eight different tissues or developmental stages. The data was taken from the International Barley Genome Sequencing Consortium (2012). A. 4-day embryos. B. Roots from seedlings (10 cm shoot stage). C. Shoots from seedlings (10 cm shoot stage). D. Young developing inflorescences (5 mm). E. Developing inflorescences (1-1.5 cm). F. Developing tillers at six-leaf stage, 3$^{rd}$ internode. G. Developing grain, bracts removed, 5 days post anthesis (DPA). H. Developing grain, bracts removed (15 DPA).

Supplemental Table 1: Markers used for genetic mapping

| Marker ID | Type | Forward sequence (5'-3') | Reverse sequence (5'-3') | Length (bp) | Enzyme | Morex* | M4205† |
|---|---|---|---|---|---|---|---|
| CAPS_2503 | CAPS | GTGAAGGCACATCACAAAGC | TGAGCAGTATGCAATCAGGG | 790 | *ScrF*I | 411, 379 | 790 |
| CAPS_2536 | CAPS | CATCACGTAAATCAAGTGCCA | TTATGCACCTCCAATGCGTA | 734 | *SfaN*I | 552, 182 | 734 |
| CAPS_2551 | CAPS | ATCATCGATCCAAGCACTCC | CAAGGCGGAGATCTGACAAT | 444 | *Ban*I | 444 | 232, 212 |
| CAPS_2558 | CAPS | ACTTCGACAAAGGACGCTGT | CTACTACCGCCAGAAGCCAC | 468 | *BstU*I | 347, 121 | 468 |
| CAPS_2560 | CAPS | ACGACGACGAAGGTACTGCT | TCTATTCCTCAGTGGCCTCG | 673 | *Sau96*I | 355, 318 | 673 |
| CAPS_2562 | CAPS | ACATCCTTGCTTGCTTGCTT | CACCCAGTGGCTCTGGTACT | 372 | *Nsi*I | 371 | 197, 175 |
| CAPS_2586 | CAPS | ATCAGAAGCCATGATGTCCC | TTGTTCTCCACACTGATGCC | 350 | *Taq*I | 195, 95, 60 | 195, 155 |
| 3_0168 | CAPS | CCCGGCTAAGTTCTGTCAAG | GACTAGGGAAACCTGCGACA | 783 | *Taq*I | 559, 224 | 389, 383 |
| Zip_2601 | CAPS | TCTTGGCCGACTCTTATTGG | AGCAGCATCTTCAGCTCCTC | 810 | *MspA1*I | 592, 218 | 366, 226, 218 |
| Zip_2613 | CAPS | ACGGTTGTAGACGGGTTGAG | ATGGGATGTGTAGGCCATGT | 854 | *AlwN*I | 625, 229 | 853 |
| Zip_2656 | CAPS | AGCAAGACTCCTCCACCAGA | CAAGGACTGTGGTGTCATGG | 797 | *Cac8*I | 436, 240, 121 | 347, 240, 121, 89 |
| Zip_2661 | CAPS | GGATCTCTTGCAGAGCCAAC | ATCGCCACAAGGTTATCAGC | 843 | *Acc*I | 843 | 660, 183 |
| Zip_2662 | CAPS | CAGCGATGCTTCGACTATCA | GATGTGATGCCCTGTTTTCC | 699 | *Mme*I | 699 | 569, 126 |
| Zip_2665 | CAPS | CAACTGGAGTGGTTGGATTG | CCTAGCCGGAAAGAAGCTC | 959 | *AlwN*I | 959 | 722, 237 |
| Zip_2667_4 | CAPS | CATGTCCCCACTGTATGATCC | GCCAAACAGGAAACACCA | 692 | *EcoR*I | 692 | 380, 312 |
| Zip_2672 | CAPS | AACGGAAACGCTCTCTCTCA | CAACCACCGCTGCTACTACA | 613 | *Fau*I | 613 | 497, 116 |
| Zip_2680_1 | CAPS | GCACTCAAGAGCAACATCCT | AGAAGGGCGGATTGTCTTCT | 850 | *Sau96*I | 777, 73 | 460, 317, 73 |
| Contig_220966 | CAPS | GCTGCCACTCTTCACAATGA | GCCGTGGAGGATCAAACTAA | 669 | *Acc*I | 518, 84, 67 | 602, 67 |
| Contig_1596897 | CAPS | TGGAGACGTATCACGCAGTC | GTACCCCTCGCCCTAAACTC | 683 | *HinP1*I | 405, 278 | 683 |
| Contig_37952_1 | CAPS | GAGTAATACCCGCCCACAAA | CGAGTCGTCCTTGAAGTTGG | 850 | *BstY*I | 485, 365 | 365, 297, 188 |
| Contig_40728_2 | CAPS | GGACCAAAGAGCCTACATGG | CCAAGAGGATGCAACTTGTG | 961 | *HinP1*I | 774, 187 | 950 |
| Contig_49785_5 | CAPS | GTTTAAGCGGGCTGTCAGAG | GTCTCATGGAGCCCAATCTC | 790 | *Hpa*II | 757, 33 | 520, 237, 33 |
| Contig_1561286_2 | CAPS | CGTGTCTGTGAAATGGTTGG | AGCGTTGTACTCCGGCTTTA | 1293 | *HinP1*I | 788, 300, 205 | 993, 300 |
| Contig_1575446 | CAPS | TGCTGCACATCTCTTTGACC | GAGAGAGCGTTTCCGTTCTG | 798 | *Nsp*I | 618, 180 | 798 |
| Contig_92279 | CAPS | CCCTAGCACATCCACCTCAT | AGACGCACACAGACGAAATG | 842 | AccI | 766,76 | 425,340,76 |

*- CAPS assay: Expected fragments size for wild-type (cv. Morex).

†- CAPS assay: Expected fragments size for Mutant (M4205).

Supplemental Table 2: Anchoring markers to the physical map of barley

| Marker ID | Morex_WGS_Contig | Chromosome | Physical Position (bp) | Corresponding BAC | FPcontig | Anchoring method |
|---|---|---|---|---|---|---|
| Contig_37952_1 | Contig_37952 | 7H | 592,633,646 | HVVMRXALLEA0035F11 | 7112 | In silico |
| Zip_2661 | Contig_65209 | 7H | 592,741,026 | HVVMRXALLHB0102L04 | 7112 | In silico |
| Zip_2662 | Contig_104939 | 7H | 592,897,645 | HVVMRX83KHA0187H24 | 7112 | In silico |
| 3_0168 | Contig_137310 | 7H | 593,142,672 | HVVMRXALLrA0254F14 | 7615 | In silico |
| Zip_2665 | Contig_263534 | 7H | 593,168,608 | HVVMRXALLmA0412B01 | 7506 | In silico |
| Zip_2667_4 | Contig_65509 | 7H | 593,203,857 | HVVMRXALLmA0412B01 | 7506 | In silico |
| Contig_1596897 | Contig_1596897 | 7H | 593,274,188 | HVVMRXALLEA0275D08 | 7506 | BAC screening |
| Zip_2672 | Contig_1575446 | 7H | 593,706,248 | HVVMRXALLEA0135I17 | 7506 | In silico/BAC screening |
| Contig_1575446 | Contig_1575446 | 7H | 593,706,248 | HVVMRXALLEA0135I17 | 7506 | In silico/BAC screening |
| Contig_49785_5 | Contig_49785 | 7H | 593,801,698 | HVVMRXALLMA0055G09 | 7506 | In silico |
| CAPS_2551 | Contig_49785 | 7H | 593,801,698 | HVVMRXALLMA0055G09 | 7506 | In silico |

Supplemental Table 3: List of the sequenced MTP BACs

| BAC ID | Chromosome | CB_start | CB_end | FPcontig | Repository | BAC assembly accession ID |
|---|---|---|---|---|---|---|
| HVVMRXALLrA0395M21 | 7H | 0 | 146 | 7112 | ENA | FJWB02025381 |
| HVVMRXALLhB0090J05 | 7H | 31 | 123 | 7112 | ENA | FJWB02001896 |
| HVVMRXALLrA0043E12 | 7H | 52 | 213 | 7112 | ENA | FJWB02002782 |
| HVVMRXALLrA0178K21 | 7H | 73 | 116 | 7112 | ENA | FJWB02025374 |
| HVVMRXALLrA0180I18 | 7H | 76 | 113 | 7112 | ENA | FJWB02002557 |
| HVVMRXALLmA0223J05 | 7H | 166 | 279 | 7112 | ENA | FJWB02003349 |
| HVVMRXALLmA0037P04 | 7H | 213 | 344 | 7112 | ENA | FJWB02003493 |
| HVVMRXALLeA0017I24 | 7H | 273 | 373 | 7112 | ENA | FJWB02005338 |
| HVVMRXALLmA0159P16 | 7H | 286 | 391 | 7112 | ENA | FJWB02025355 |
| HVVMRXALLeA0098N12 | 7H | 359 | 426 | 7112 | ENA | FJWB02025330 |
| HVVMRXALLhA0023E08 | 7H | 393 | 486 | 7112 | ENA | FJWB02025341 |
| HVVMRXALLrA0254F14 | 7H | 0 | 134 | 7615 | ENA | FJWB02025378 |
| HVVMRXALLeA0172A05 | 7H | 53 | 134 | 7615 | ENA | FJWB02044216 |
| HVVMRXALLmA0220E09 | 7H | 22 | 123 | 7615 | ENA | FJWB02039734 |
| HVVMRXALLmA0230A06 | 7H | 62 | 159 | 7615 | ENA | FJWB02039814 |
| HVVMRXALLhA0032I06 | 7H | 63 | 169 | 7615 | ENA | FJWB02025342 |
| HVVMRXALLhB0145H15 | 7H | 0 | 99 | 4483 | ENA | FJWB02006610 |
| HVVMRXALLhB0138E15 | 7H | 45 | 142 | 4483 | ENA | FJWB02006575 |
| HVVMRXALLrA0045N01 | 7H | 78 | 156 | 4483 | ENA | FJWB02006662 |
| HVVMRXALLmA0337G17 | 7H | 109 | 211 | 4483 | ENA | FJWB02000707 |
| HVVMRXALLmA0155C21 | 7H | 174 | 289 | 4483 | ENA | FJWB02002639 |
| HVVMRXALLmA0299O06 | 7H | 258 | 382 | 4483 | ENA | FJWB02005902 |
| HVVMRXALLmA0465P05 | 7H | 312 | 443 | 4483 | ENA | FJWB02004379 |
| HVVMRXALLmA0510H04 | 7H | 378 | 508 | 4483 | ENA | FJWB02001578 |
| HVVMRXALLmA0381J10 | 7H | 425 | 556 | 4483 | ENA | FJWB02007510 |
| HVVMRXALLeA0143C22 | 7H | 496 | 590 | 4483 | ENA | FJWB02003880 |
| HVVMRXALLrA0400I23 | 7H | 0 | 133 | 44845 | ENA | FJWB02025382 |
| HVVMRXALLrA0226A20 | 7H | 27 | 114 | 44845 | ENA | FJWB02001332 |
| HVVMRXALLeA0131L13 | 7H | 77 | 157 | 44845 | ENA | FJWB02005172 |
| HVVMRXALLhC0124B19 | 7H | 128 | 211 | 44845 | ENA | FJWB02000301 |
| HVVMRXALLeA0089H17 | 7H | 130 | 235 | 44845 | ENA | FJWB02006334 |
| HVVMRXALLmA0510A15 | 7H | 194 | 285 | 44845 | ENA | FJWB02001577 |
| HVVMRXALLmA0037G09 | 7H | 223 | 351 | 44845 | ENA | FJWB02003485 |
| HVVMRXALLmA0169C16 | 7H | 225 | 303 | 44845 | ENA | FJWB02000810 |
| HVVMRXALLmA0089A11 | 7H | 339 | 442 | 44845 | ENA | FJWB02006485 |
| HVVMRXALLhB0142P06 | 7H | 398 | 471 | 44845 | ENA | FJWB02006596 |
| HVVMRXALLmA0272A07 | 7H | 432 | 532 | 44845 | ENA | FJWB02002497 |
| HVVMRXALLmA0180L03 | 7H | 489 | 601 | 44845 | ENA | FJWB02000870 |
| HVVMRXALLmA0416P19 | 7H | 554 | 692 | 44845 | ENA | FJWB02025362 |
| HVVMRX83KhA0095B11 | 7H | 621 | 723 | 44845 | ENA | FJWB02004898 |
| HVVMRXALLmA0457P05 | 7H | 679 | 776 | 44845 | ENA | FJWB02003818 |
| HVVMRXALLhB0062D15 | 7H | 712 | 794 | 44845 | ENA | FJWB02003700 |
| HVVMRXALLmA0268L16 | 7H | 744 | 865 | 44845 | ENA | FJWB02002468 |
| HVVMRXALLeA0192I19 | 7H | 810 | 933 | 44845 | ENA | FJWB02006221 |
| HVVMRXALLhB0069D14 | 7H | 849 | 1048 | 44845 | ENA | FJWB02001812 |
| HVVMRXALLhB0103F17 | 7H | 0 | 114 | 7506 | ENA | FJWB02005652 |
| HVVMRXALLhA0054F13 | 7H | 1 | 56 | 7506 | ENA | FJWB02001430 |
| HVVMRXALLmA0125F11 | 7H | 66 | 171 | 7506 | ENA | FJWB02007923 |
| HVVMRXALLeA0272A06 | 7H | 79 | 172 | 7506 | ENA | FJWB02025335 |
| HVVMRXALLmA0208I02 | 7H | 124 | 226 | 7506 | ENA | FJWB02004775 |
| HVVMRXALLhA0037N16 | 7H | 163 | 227 | 7506 | ENA | FJWB02001407 |
| HVVMRXALLmA0258A21 | 7H | 197 | 264 | 7506 | ENA | FJWB02002905 |
| HVVMRXALLeA0213D22 | 7H | 223 | 328 | 7506 | ENA | FJWB02002042 |
| HVVMRXALLmA0340A18 | 7H | 251 | 365 | 7506 | ENA | FJWB02000716 |
| HVVMRXALLeA0135I17 | 7H | 307 | 417 | 7506 | ENA | FJWB02005201 |
| HVVMRXALLmA0477O22 | 7H | 352 | 463 | 7506 | ENA | FJWB02004413 |
| HVVMRXALLeA0275D08 | 7H | 415 | 522 | 7506 | ENA | FJWB02005800 |
| HVVMRXALLrA0380O16 | 7H | 438 | 469 | 7506 | ENA | FJWB02005042 |
| HVVMRXALLeA0188J13 | 7H | 464 | 539 | 7506 | ENA | FJWB02006179 |
| HVVMRXALLmA0412B01 | 7H | 465 | 566 | 7506 | ENA | FJWB02025361 |

Supplemental Table 4: Primers used in this study

| Primer ID | Forward sequence (5'-3') | Reverse sequence (5'-3') | Length (bp)* |
|---|---|---|---|
| I. Primers used for TILLING analysis | | | |
| AK366098_Exon1 | GGGTCCAGATTGATTCATCC | GCAGTGCAGGCATTTCAATC | 1376 |
| AK366098_Exon3 | ATCAGGGAGCATGGTTTACG | AGCCGTCATCTGCTTCACTT | 596 |
| II. Primers used for subcellular localization experiments | | | |
| pSB179 | CAGACGGGATCGATCTAGGA | GAACTTCAGGGTCAGCTTGC | N.A. |
| HvAs_SC_WT | TCACTAGTATGGCGTCGTCCTGCATCCCGACGG | CTAAGCTTTGGCCTCTTTCTCGCCGGCTTTCTGC | 1394 |
| HvAs_SC_M4205 | TCACTAGTATGGCGTCGTCCTGCATCCCGACGG | CAAAGCTTTGAAACAGTTCAATATCTGAAAGTTTA | 1193 |
| HvAs_SC_TILLING | TCACTAGTATGGCGTCGTCCTGCATCCCGACGG | CTAAGCTTTAGTCTTCTTCTTTTTCTTCTCCACC | 944 |
| HvAs_SC_cTP | TCACTAGTATGGCGTCGTCCTGCATCCCGACGG | CTAAGCTTTGGCGAGCAGCGCGGCCGCCTCGTT | 266 |
| HvAs_SC_CCT | TCACTAGTATGAGGGAAGGCAGCGTACAGAAATTGA | CAAAGCTTCCGCCTGGCTGACAAACCTCCCCTTG | 158 |
| III. Primers used for analysis of transgenic lines derived from site-directed mutagenesis | | | |
| HYG | CATGGTGGAGCACGACACTCTC | GATCGGACGATTGCGTCGCA | 1567 |
| Cas9 | TTTAGCCCTGCCTTCATACG | TTAATCATGTGGGCCAGAGC | 734 |
| OsU3 | CAGGGACCATAGCACAAGAC | TCAGCGGGTCACCAGTGTTG | 595 |
| Target Motif 1 | GGCGCTCAAGGCGTGGTATGACAG | TCAGCGGGTCACCAGTGTTG | N.A |
| Target Motif 2 | GGCGCTGATGCGTCATCCACTGAC | TCAGCGGGTCACCAGTGTTG | N.A |

* N.A. - Not applicable.

| Supplemental Table 5: Identified TILLING mutants for *HvAST* | | | | | | |
|---|---|---|---|---|---|---|
| Plant family ID* | SNP position† | SNP | Orignal‡ | $M_2/M_4$ status | Effect | Region |
| 10580-1 | 30 | C/T | AT**C** | Heterozygote | Ile/Ile | Exon 1 |
| 3283-1 | 60 | G/C | GC**G** | Heterozygote | Ala/Ala | Exon 1 |
| 12436-1 | 72 | C/T | GC**C** | Heterozygote | Ala/Ala | Exon 1 |
| 9688-1 | 114 | G/A | CC**G** | Homozygote | Pro/Pro | Exon 1 |
| 3227-1 | 130-138 | Deletion | **TCCTCGGCG** | Homozygote | Ser Ser Ala/--- | Exon 1 |
| 7924-1 | 156 | C/T | AA**C** | Homozygote | Asn/Asn | Exon 1 |
| 9414-1 | 168 | C/T | GC**C** | Heterozygote | Ala/Ala | Exon 1 |
| 14946-1 | 222 | C/T | AC**C** | Heterozygote | Thr/Thr | Exon 1 |
| 9878-1 | 279 | G/T | GG**G** | Homozygote | Gly/Gly | Exon 1 |
| 8265-1 | 286 | G/A | **G**CG | Homozygote | Ala/Thr | Exon 1 |
| 6646-1_24_1 | 286 | C/T | GA**C** | Homozygote | Asp/Asp | Exon 1 |
| 9825-1 | 286 | C/T | **C**TC | Homozygote | Leu/Phe | Exon 1 |
| 13029-1 | 367 | C/T | **C**CC | Heterozygote | Pro/Ser | Exon 1 |
| 10459-1 | 367 | C/T | CT**C** | Homozygote | Leu/Leu | Exon 1 |
| 3779-1 | 427 | C/T | **C**CG | Homozygote | Pro/Ser | Exon 1 |
| 12772-1 | 435 | C/T | AG**C** | Homozygote | Ser/Ser | Exon 1 |
| 13269-1 | 504 | C/T | TC**C** | Heterozygote | Ser/Ser | Exon 1 |
| 10799-1 | 514 | C/T | **C**CG | Homozygote | Pro/Ser | Exon 1 |
| 9920-1 | 634 | G/A | **G**CC | Homozygote | Ala/Thr | Exon 1 |
| 13239-1 | 641 | G/A | G**G**G | Heterozygote | Gly/Glu | Exon 1 |
| 2889-1 | 642 | G/A | GG**G** | Heterozygote | Gly/Gly | Exon 1 |
| 2920-1 | 659 | G/A | G**G**C | Heterozygote | Gly/Asp | Exon 1 |
| 7669-1 | 664 | C/T | **C**TC | Homozygote | Leu/Phe | Exon 1 |
| 3698-1 | 667 | A/T | **A**GT | Homozygote | Ser/Cys | Exon 1 |
| 6912-1 | 695 | C/T | A**C**T | Heterozygote | Thr/Ile | Exon 1 |
| 12113-1 | 730 | C/T | **C**CC | Heterozygote | Pro/Ser | Exon 1 |
| 8222-1 | 774 | C/T | CA**C** | Heterozygote | His/His | Exon 1 |
| 12189-1 | 821 | G/A | A**G**C | Heterozygote | Ser/Asn | Exon 1 |
| 4394-1 | 847 | C/T | **C**CA | Heterozygote | Pro/Ser | Exon 1 |
| 12710-1 | 886 | G/A | **G**CG | Homozygote | Ala/Thr | Exon 1 |
| 10022-1 | 909 | G/A | GA**G** | Homozygote | Glu/Glu | Exon 1 |
| 6460-1 | 928 | A/T | **A**AG | Heterozygote | Lys/stop codon | Exon 1 |
| 10996-1 | 1009 | G/A | **G**AT | Heterozygote | Asp/Asn | Exon 1 |
| 13603-1 | 1009 | G/A | **G**AT | Heterozygote | Asp/Asn | Exon 1 |
| 10972-1 | 1032 | G/A | AA**G** | Heterozygote | Lys/Lys | Exon 1 |
| 3812-1 | 1062 | C/T | CT**C** | Heterozygote | Leu/Leu | Exon 1 |
| 11403-1 | 2358 | G/A | A**G**C | Homozygote | Ser/Asn | Exon 2 |
| 13059-1 | 2379 | G/A | AA**G** | Homozygote | Lys/Lys | Exon 2 |
| 14134-1 | 2385 | G/A | AA**G** | Heterozygote | Lys/Lys | Exon 2 |
| 3005-1 | 2430 | G/A | CG**G** | Heterozygote | Arg/Arg | Exon 2 |
| 4217-1 | 2434 | G/A | **G**TG | Homozygote | Val/Met | Exon 2 |
| 11797-1 | 2441 | C/T | G**C**C | Heterozygote | Ala/Val | Exon 2 |

* Plant identifers are referring to $M_2$ TILLING family with one exception 6646-1_24_1 in regarding to $M_4$.

† Coordinates based on genomic sequence of cv. Barke. The Adenine of start codon is counted as position +1.

‡ The SNP position is marked in bold.
